# Bistable perception of symbolic numbers

**DOI:** 10.1101/2024.06.04.597279

**Authors:** Junxiang Luo, Isao Yokoi, Serge O. Dumoulin, Hiromasa Takemura

**Author notes:** **Correspondence** Junxiang Luo, Hiromasa Takemura.

## Abstract

Numerals, i.e., semantic expressions of numbers, enable us to have an exact representation of the amount of things. Visual processing of numerals plays an indispensable role in the recognition and interpretation of numbers. Here, we investigate how visual information from numerals is processed to achieve semantic understanding. We first found that partial occlusion of some digital numerals introduces bistable interpretations. Next, by using the visual adaptation method, we investigated the origin of this bistability in human participants. We showed that adaptation to digital and normal Arabic numerals, as well as homologous shapes, but not Chinese numerals, biases the interpretation of a partially occluded digital numeral. We suggest that this bistable interpretation is driven by intermediate shape processing stages of vision, i.e., by features more complex than local visual orientations but more basic than the abstract concepts of numerals.

## Introduction

The sense of number endows animals with a better chance of survival by bringing behavioral advantages to daily activities like foraging and avoiding predation (Nieder 2020). Psychophysical and neuroscientific studies using arrays of dots demonstrated that the neural mechanisms for processing non-symbolic number information are shared among humans and other animals (Burr & Ross 2008, Burr et al 2017, Harvey et al 2013, Hubner & Schutz 2017, Nieder 2012, Nieder 2013, Nieder 2016, Park et al 2016, Tsouli et al 2022, Viswanathan & Nieder 2013).

However, one aspect that dissociates humans from other animals is the ability to recognize symbolic numbers. The semantic understanding of those characters representing numbers, i.e., numerals, enables humans to have a much more exact and flexible appreciation of the numerical world (Wiese 2003), and also mathematical and logical skills that lead to significant developments in sciences, technologies, and social economics.

In the human brain, patches on the inferior temporal gyrus represent symbolic numbers, such as Arabic numerals (Cai et al 2023, Conrad et al 2023, Shum et al 2013, Yeo et al 2020, Yeo et al 2017). However, as of yet, we do not know how the recognition of symbolic numbers is generated through visual processing from local orientation to complex shapes, category information, and semantic recognition (Kravitz et al 2013, Riesenhuber & Poggio 1999).

Bistable perception is a powerful tool for investigating the relationship between perception and visual processing (Blake & Logothetis 2002, Brascamp & Shevell 2021, Leopold & Logothetis 1999, Rodriguez-Martinez & Castillo-Parra 2018, Schneider et al 2019, Sterzer et al 2009). Specifically, it enables scientists to examine what factors lead to a specific interpretation while the physical input to the visual system remains identical.

Here, we introduce a bistable perception of symbolic numbers by partial occlusion. We used digital numerals that widely exist in modern society. Because of their simple design, they are applied to various consumer electronics, such as calculators and traffic countdown timers (***Fig. 1A***). The semantic interpretation of digital numerals can be easily manipulated by adding or removing one stick from the character, such as those shown in ***Fig. 1B***, where the interpretation is altered between 5 and 9, and 6 and 8. Inspired by this feature, we found that a partial occlusion of these symbolic characters results in bistable semantic interpretations. Such as the case in ***Fig. 1C***, people recognize this occluded digital numeral as either 6 or 8. This phenomenon can also be extended to a group of sequentially arranged numerical characters, the partial occlusion of them leads to multiple interpretations which occur most noticeably in monotonically increasing and decreasing orders (***Fig.1D***). This bistable interpretation of digital numerals is also present in our daily life. ***Supplementary Video 1*** demonstrates an example of a traffic countdown timer in the city. When part of the numerical character is occluded by the protective cover, starting from one moment, people tend to see the numbers being counted from 5 to 9 instead of being counted from 9 to 5.

**Figure 1.**
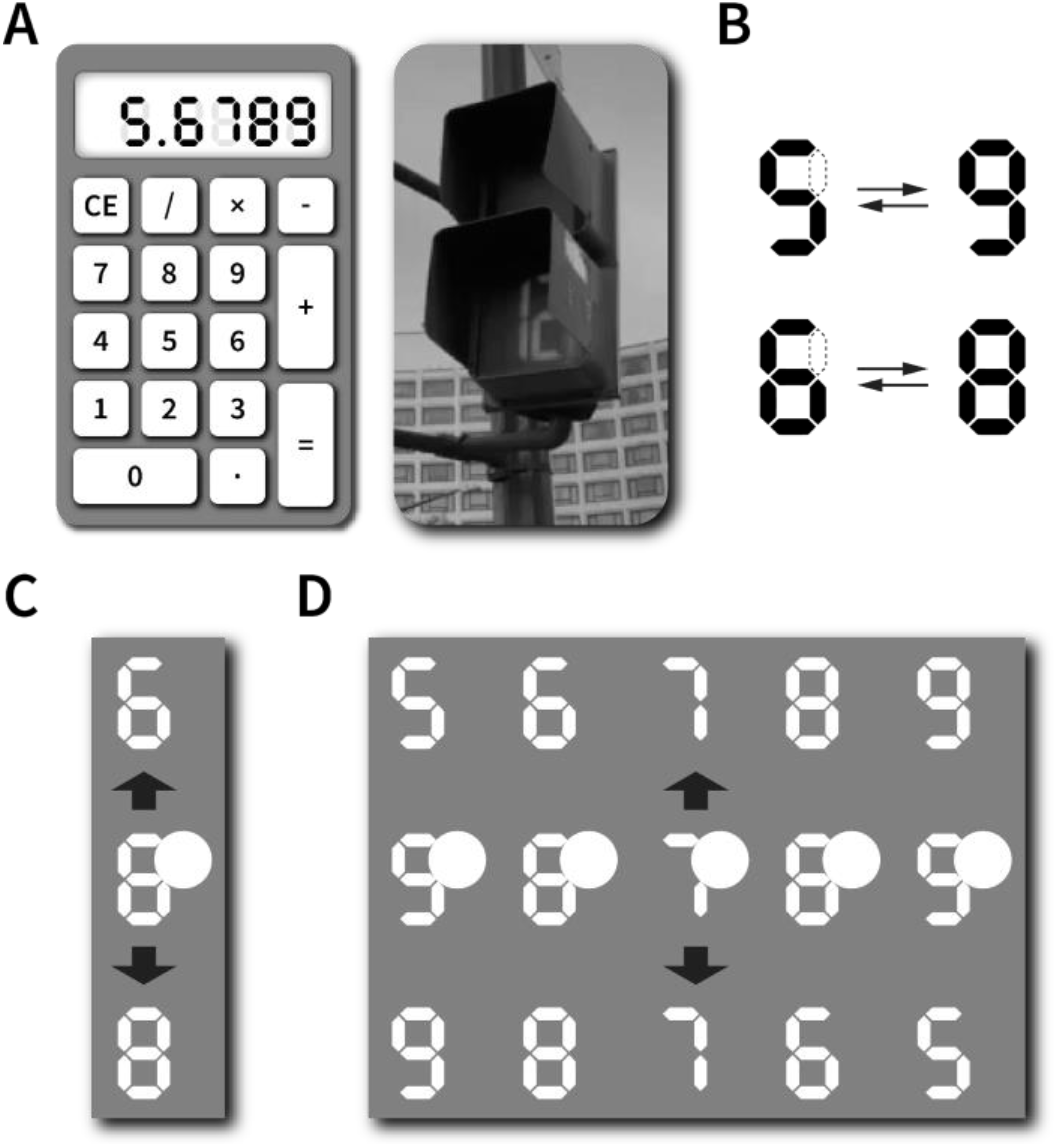
Digital numerals and perceptual bistability. (**A**) Digital numerals are shown in the display of a calculator (*left*) and traffic countdown timer (*right*). (**B**) Adding or removing the top-right stick of some digital numerals changes their semantic interpretations. (**C**) The middle digital numeral is partially masked by a disk of the same luminance and therefore can be interpreted as either 6 (*top character*) or 8 *(bottom character*). (**D**) The manipulation of occlusion that is applied to a group of numerical characters (*middle row*) causes two different interpretations of ordered sequences, e.g., they can be interpreted as an ascending (*top row*) or descending (*bottom row*). order

In this study, we aimed to shed light on the mechanisms underlying the visual processing of symbolic numbers by psychophysically measuring and manipulating the bistable perception of occluded digital numerals.

Adaptation is an effective psychophysical method to selectively manipulate the sensitivity of the visual system and target the cortical loci that contribute to visual functions (Frisby 1979, Kohn 2007, Webster 2015). For example, if there were clusters of neurons selectively encoding a given numeral (such as 6), prolonged exposure to the visual presentation of that numeral will reduce the responsiveness of those neurons and potentially influence the perceptual recognition of that number. Therefore, testing how adaptation impacts the perception of partially occluded digital numerals should reveal the perceptual and neural mechanisms underlying the visual processing of numerals.

In this study, we employed a psychophysical task using several representative symbols and shapes as adaptation stimuli. We assessed whether we could bias the semantic recognition of occluded digital numerals (***Fig. 2***). We found that adaptation to stimuli that shared certain shape similarities with the partially occluded digital numerals resulted in a strong perceptual bias. In contrast, other adaptation stimuli with the same semantic meaning but different shapes (Chinese numerals) did not have such an effect. Results indicate that the perceptual bistability of digital numerals have a neural origin, which is involved in processing complex visual form information.

**Figure 2.**
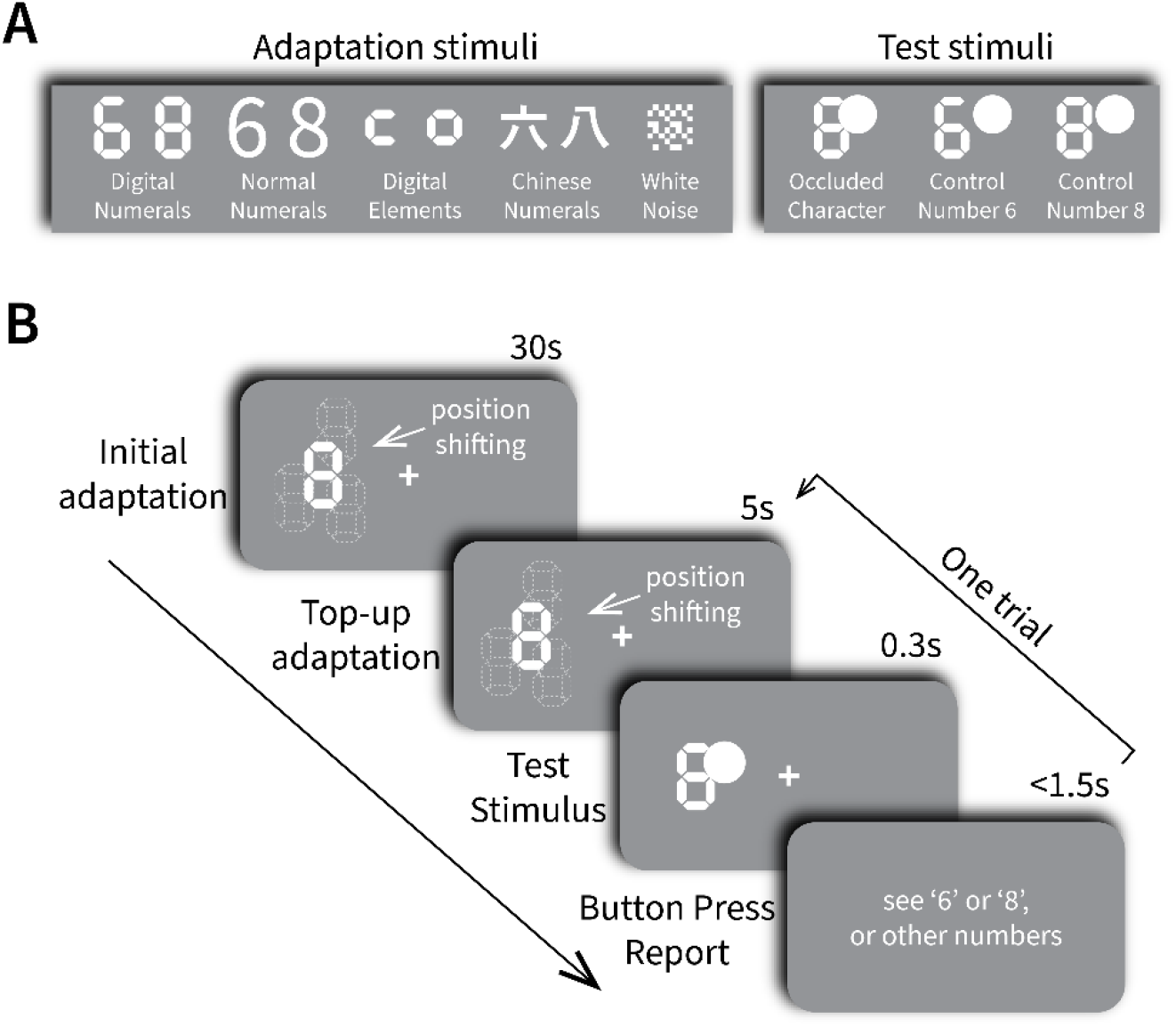
Illustrations of stimuli designs and task time series. (**A**) Five types of visual stimuli were used as adaptation stimuli, including two kinds of numerical characters (digital and normal Arabic numerals), one kind of shape stimulus (digital elements), one kind of Chinese numerals (semantically representing 6 and 8), and control stimulus (white noise). Three characters were presented as test stimuli after adaptation, including one occluded character (with a mask covering the right-upper part), and two control characters (with a mask on the right-upper side, but no coverage). (**B**) Each experimental block started with an initial adaptation period where normal numerical characters or shapes were presented for 30 seconds. It was followed by a loop of trials. Each trial sequentially contains a top-up (5 seconds) adaptation to the same adaptation stimuli as used in the initial adaptation, a presentation period for the test stimulus, and a report time window, participants were asked to report whether they recognize the occluded numeral as 6 or 8, or other numbers, by pressing one of three buttons on a gaming pad.

## Methods

### Participants

Seventeen adults (9 females and 8 males; ages ranging from 19 to 31 years) with normal or corrected-to-normal visual acuity participated in the experiment. Tasks were performed after they signed a written informed consent which included explanations of voluntary participation, experimental safety, and freedom to withdraw, as well as an agreement on the data sharing policy. All participants were born in Japan and were proficient in Japanese and in reading Chinese characters. All participants passed the battery of tests assessing visual functions including acuity, astigmatism, and binocular stereopsis before the main experiment. The study protocol was approved by the Ethics Committee of National Institutes of Natural Sciences (protocol number: EC01-64). We note that the data from one participant were excluded from analyses due to excessive eye movements (see below). Therefore, we report the data acquired from 16 participants (8 females and 8 males; ages ranging from 19 to 31 years).

### Apparatus

Visual stimuli were presented on a BenQ 24.5-inch display (ZOWIE XL2546K, BenQ, Taipei) with a spatial resolution of 1920×1080 pixels and a refresh rate of 240 Hz. The luminance of the display was linearly scaled using gamma calibration. A self-designed chinrest mounted 65 cm away from the display was used to stabilize participants’ heads and to keep the observation distance constant. Fixation was monitored by an eye-tracking system (LiveTrack Lightning, Cambridge Research Systems, Rochester, United Kingdom); the camera was placed in front of the chinrest and 18 cm away from the eyes and tracked the positions of both eyes online. Visual stimuli generation and online data analysis were performed using a workstation (Dell Precision 3650, Dell, Round Rock, TX, USA) operated under the Linux system (Ubuntu 20.04 LTX). Participants used a gaming pad (Retro-Bit Legacy 16, Retro-Bit, Pomona, CA, USA) to report their perceptions.

### Stimuli

Visual stimuli were designed using MATLAB Psychtoolbox extensions (Brainard 1997, Kleiner et al 2007; http://psychtoolbox.org/, Pelli 1997). All stimuli were drawn in white and presented against a medium-gray background. During stimulus presentation, a white fixation cross of 0.5 degrees in diameter was also presented at the center of the display (***Fig. 2B***).

We used five different types of adaptation stimuli (***Fig. 2A left***): (1) digital numerals 6 and 8, (2) normal Arabic numerals 6 and 8, (3) two digital elements representing the upper part of digital numerals 6 and 8, (4) Chinese numerals 6 and 8 (六 and 八 corresponding to 6 and 8), and (5) white noise patterns as control stimuli. There were two white noise stimuli, generated by random shuffling of the pixels in the two digital numerals. Since we did not find any significant difference in behavioral responses between the two white noise stimuli, we present the averaged results of the two white noise stimuli in the population analysis. The height of the adaptation stimuli was 200 pixels (corresponding to a visual angle of about 5 degrees, the height was halved in the digital element stimuli). The width of the digital and normal Arabic numerals, the digital elements, and the white noise pattern was 124 pixels (3.3 deg), while the width of the Chinese numerals was 216 pixels (5.8 deg) because of their specific design.

We used three test stimuli (***Fig. 2A right***). The first one was a digital numeral with a white circular mask placed over the right upper part. The second and third ones were control test stimuli. Each control test stimulus was composed of either the digital numeral 6 or 8 with the same mask but without overlap, so that the digital numerals could be easily recognized as either 6 or 8 without ambiguity. The test digital numerals were of the same size as the adaptation digital numerals, whereas the diameter of the mask was 178 pixels (4.8 deg). The control test stimuli were used to make sure that participants were always making valid responses during the task. Since all participants responded correctly to the control test stimuli in 100% of the trials, we considered their perceptual responses to the partially occluded stimuli also reliable.

***Figure 2B*** depicts the time series of the experiment. Both adaptation and test stimuli were presented in either the left or right visual field. The adaptation stimuli were presented horizontally and on average 6 degrees away from the fixation cross, while their positions were randomly shifted every second within a circular area of 6 degrees. Such position shift was used to prevent low-level adaptation effects in early visual areas (e.g., the primary visual cortex) where the neurons have smaller receptive field sizes, and also give enough distance from the fixation point to ensure that they did not fall onto the fovea. The test stimuli were presented at an eccentricity of 6 deg but were kept static during presentation.

Both adaptation and test stimuli were shown on either the left or right side of the visual field within one task block, but their positions were changed to the other side in the next task block.

### Procedure

During the experiment, participants were asked to maintain fixation at the central white cross, while paying attention to the peripheral visual stimuli. The initial adaptation period lasted for 30 seconds before the loop of trials (see ***Supplementary Video 2***). In each trial, a 5-second top-up adaptation to the same stimulus was followed by a 300-millisecond presentation of the test stimulus. At the end of each trial, participants were instructed to report whether they recognize the test stimulus as 6 or 8, or other numbers, by pressing the ‘L’ or ‘R’, or ‘A’ button on the gaming pad (***Figure 2B***). The experiment was divided into five sessions based on five different adaptation stimuli as shown in ***Fig. 2A left***. There were four blocks in each session; in two blocks, the adaptation stimulus was number 6 or the element corresponding to number 6; and in the other two blocks, number 8 or the element corresponding to number 8. In both pairs of blocks, stimuli were presented on either the left or right visual field in one block and the opposite visual field in the other block. Within each block, every combination of the adaptation stimulus and the three test stimuli (***Fig. 2A right***) was repeated 10 times (adaptation to white noise was repeated five times). So, there were a total of 20 repetitions for each combination (10 repetitions for white noise and test stimuli combination). As a result, there were 540 trials in the whole experiment. Each block consisting of 30 trials lasted around five minutes (15 trials and 2.5 minutes for the white noise adaptation condition), resulting in a 90-minute experiment.

### Data analysis and quantification

The proportions of trials in which participants recognized the occluded digital numeral as either 6, 8, or other numbers were calculated for each adaptation condition. We quantified a perceptual bias in each condition by calculating the bias index using the following equation:

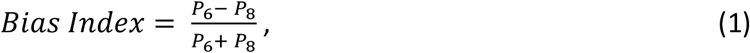

where *P*_*6*_ and *P*_*8*_ indicate the proportions of trials in which participants reported “6” and “8”, respectively. The bias index ranged from -1 to 1; positive values indicate that participants reported recognizing the occluded numeral as 6 more often than 8 while negative values indicate that participants reported 8 more often than 6. The population distributions of the bias index across 16 participants are illustrated using a violin plot (https://zenodo.org/records/4559847) in ***Figure 3***, where the shadow represents a kernel density estimate of the data. The statistical significance of the difference in bias indices between adaptation conditions was tested by the Krusukal-Wallis one-way analysis of variance. We also calculated Cohen’s d to measure the effect size. To further evaluate the weight of perceptual bias differences between adaptation to numbers 6 and 8, we performed the Bayesian Wilcoxon signed-rank test and calculated Bayes factors (BF_10_) for each adaptation condition using the JASP statistical software (https://jasp-stats.org/; JASP Team, 2024). A larger BF_10_ indicates stronger evidence for different perceptual biases between adaptation to numbers 6 and 8, while a smaller BF_10_ indicates evidence for the same perceptual biases.

**Figure 3.**
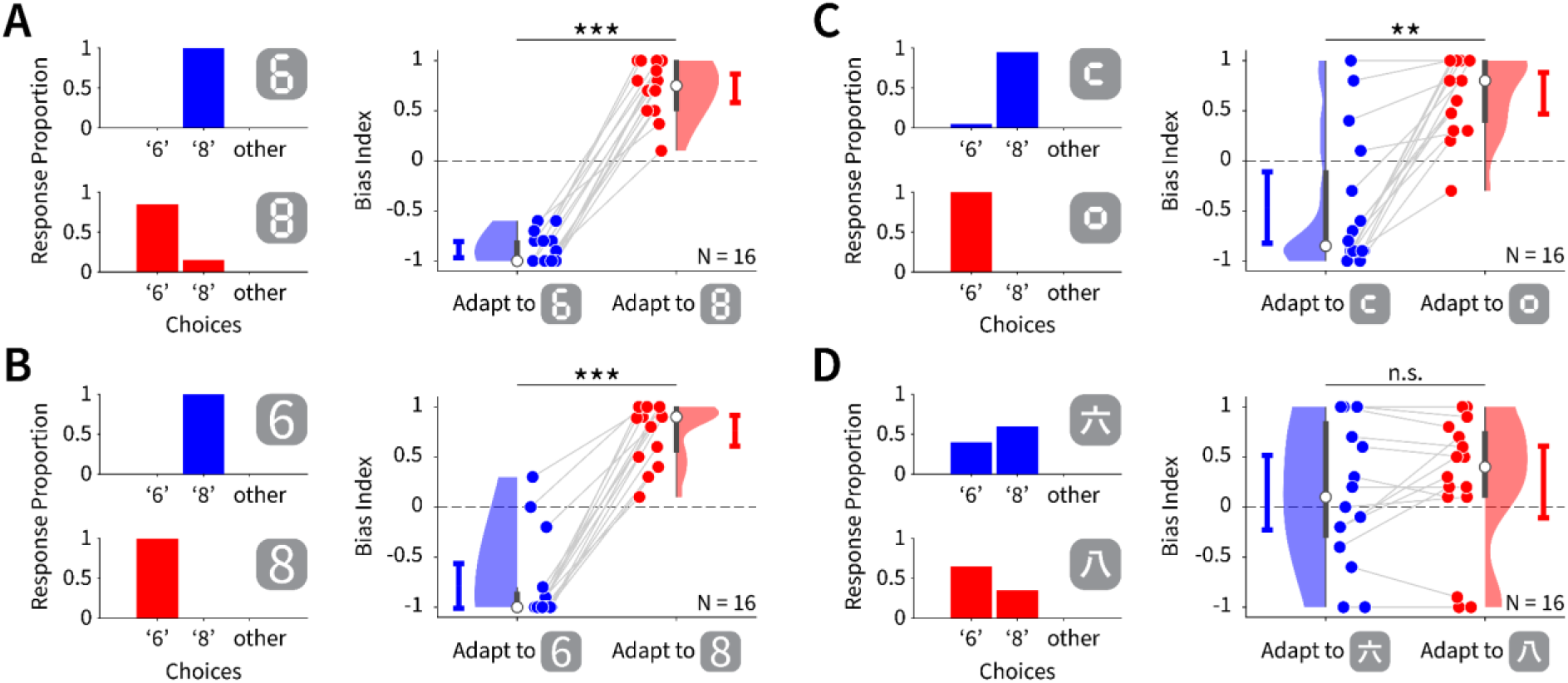
Single participant and populational results of four adaptation conditions. (**A**) *Left*: response proportions of one participant (P1) reporting seeing 6, 8, or other numbers, after adaptation to digital numeral 6 (*upper panel*) and digital numeral 8 (*lower panel*). The results show strong but opposite perceptual biases in two adaptation conditions. *Right*: populational results of the same adaptation conditions from 16 participants. Similar to the results of a single participant, bias indices of two conditions distribute oppositely and are significantly different from each other (blue: adaptation to digital numeral 6; red: adaptation to digital numeral 8). Dots depict the data from individual participants. Dots of different conditions but the same participant are connected by gray lines. Shadows represent a kernel density estimate of the data. The error bar depicts a 95% confidence interval; (**B**) Same description as in (**A**), but the conditions are adaptation to normal Arabic numerals 6 and 8; (**C**) Same description as in (**A**), but the conditions are adaptation to digital element ㄷ and ㅁ; (**D**) Same description as in (**A**), but the conditions are adaptation to Chinese numerals 六and 八, there is no significant difference between two conditions. ***: *p*<0.001; **: *p*<0.01; n.s.: not significant in Krusukal-Wallis one-way analysis of variance.

The eye fixation stability of each participant was monitored to determine whether their data were included in the statistical analysis. Data from one participant were excluded from the analysis due to an excessive amount of saccadic eye movement to the adaptation stimuli.

### Code availability

Data and code for visual stimuli presentation as well as data-analysis are available in a public repository and will become open to the public after an acceptance of the manuscript.

## Results

We first tested whether adaptation to the digital numerals 6 and 8 (***Fig. 2A Left***), which share the same shape as the test stimuli (***Fig. 2B Right***), biases the recognition of the partially occluded digital numeral. Results from one representative participant (P1) are illustrated in ***Figure 3A Left***. After adaptation to digital numerals, the responses to the partially occluded digital numeral showed clear perceptual biases despite the ambiguity caused by the partial occlusion. Specifically, after adaptation to digital numeral 6, the participant always reported the test stimulus as number 8 (***Fig. 3A Left top***, response proportions: 0% for 6; 100% for 8, and 0% for other numbers); whereas after adaptation to digital numeral 8, the participant reported the test stimulus as number 6 in most trials (***Fig. 3A Left bottom***, response proportions: 85% for 6, 15% for 8, and 0% for other numbers).

This perceptual bias was confirmed at the population level (***Fig. 3A Right***). The bias indices of all 16 participants were negative under the adaptation condition of digital numeral 6 (*mean* = -0.89; 95% confidence interval [*CI*]: ± 0.08), indicating that participants tended to see the test stimulus as 8. On the other hand, the bias indices were all positive under the adaptation condition of digital numeral 8 (*mean* = 0.72 ± 0.14 [95% *CI*]), indicating participants tended to recognize the test stimulus as 6. The perceptual biases were significantly different between the two adaptation conditions (*d* = 7.00; *p* = 3.45 × 10^-6^; *BF*_*10*_ = 1.1 × 10^4^). These results show that the recognition of partially occluded digital numerals was strongly biased by visual adaptation. This effect of adaptation suggests that the bistable perception is not driven by the subjective interpretation of the semantics of numerals, but constitutes a visual information processing mechanism.

Next, we asked which level of visual processing determines the bistable perception of the occluded digital numeral. We note that the position shifting of adaptation stimuli in our task paradigm already suggests that the perceptual biases do not happen in the early stages of visual processing (such as the primary visual cortex), where the receptive fields of neurons are relatively small and the adaptation effect is strictly localized (Kohn 2007, Kohn & Movshon 2003).

To investigate whether the later stages of visual processing play a role in bistable perception, we introduced normal Arabic numerals as adaptation stimuli (***Fig. 2A Left***). Normal Arabic numerals and digital numerals have similar global appearances, but differ in their local visual features; digital numerals are comprised of horizontally and vertically oriented intermittent straight sticks, whereas normal Arabic numerals are comprised of continuous curvatures. Since primary visual processing stages are sensitive to local feature differences whereas later stages are not (Kravitz et al 2013), examining whether adaptation to the normal Arabic numerals influence the recognition of the occluded digital numeral allowed us to answer whether later stages of visual processing played a fundamental role in the perceptual bistability of digital numerals. For example, compared with early visual areas, intermediate visual areas such as V4, are driven less by local features of the stimulus (e.g. edges) but rather by more complex features (e.g. curvature; Dumoulin & Hess 2007, Gallant et al 1993, Pasupathy & Connor 2002, Wilkinson et al 2000, Wilson et al 1997).

Results showed that adaptation to normal Arabic numerals is comparable with adaptation to digital numerals, both at the single participant level (***Fig. 3B Left***, response proportions of adaptation to normal Arabic numeral 6: 0% for 6, 100% for 8, and 0% for other numbers; response proportions of adaptation to normal Arabic numeral 8: 100% for 6, 0% for 8, and 0% for other numbers) and at the population level (***Fig. 3B Right***, bias indices of adaptation to normal Arabic numeral 6, *mean* = -0.79 ± 0.22 [95% *CI*]; bias indices of adaptation to normal Arabic numeral 8, *mean* = 0.76 ± 0.15 [95% *CI*]; *d* = 4.07, *p* = 9.77 × 10^-7^; *BF*_*10*_ = 2.8 × 10^3^). The adaptation effects of normal Arabic numerals further support the idea that bistable recognition of the occluded digital numerals is established later in the visual processing stream, rather than in the early stages.

Lastly, we investigated whether the processing of semantics plays a critical role in generating the bistable perception. We first tested the effect of adaptation to digital elements (the upper parts of the digital numerals, ***Fig. 2A Left***); this modification removed the semantic components of the digital numerals while preserving some shape similarities. The results were similar to those in digital and normal Arabic numerals adaptation conditions at both the single participant level (***Fig. 3C Left***, response proportions of adaptation to digital element ㄷ: 5% for 6, 95% for 8, and 0% for other numbers; response proportions of adaptation to digital element ㅁ: 100% for 6, 0% for 8, and 0% for other numbers) and population level (***Fig. 3C Right***, bias indices of adaptation to digital element ㄷ, *mean* = -0.47 ± 0.36 [95% *CI*]; bias indices of adaptation to digital element ㅁ, *mean* = 0.67 ± 0.21 [95% *CI*]; *d* = 1.98, *p* = 0.0065; *BF*_*10*_ = 1.1 × 10^3^), although the effect size of differences between two adaptation conditions was lower than that obtained under the adaptation conditions of digital and normal Arabic numerals.

As opposed to digital elements, Chinese numerals share the same meanings as digital numerals, while their shapes are different (***Fig. 2A Left***). We predicted that if the semantic processing stage plays a role in the bistable perception, there would be perceptual bias after adapting to Chinese numerals. Results showed that adaptation to Chinese numerals did not bias perceptual interpretation of the occluded digital numerals at either the single participant level (***Fig. 3D Left***, response proportions of adaptation to 六: 40% for 6, 60% for 8, and 0% for other numbers; response proportions of adaptation to 八: 65% for 6, 35% for 8, and 0% for other numbers) or the population level (***Fig. 3D Right***, bias indices of adaptation to Chinese numeral 六, *mean* = 0.14 ± 0.37 [95% *CI*]; bias indices of adaptation to Chinese numeral 八, *mean* = 0.25 ± 0.36 [95% *CI*]; *d* = 0.15, *p* = 1.00, *BF*_*10*_ = 0.31). Moreover, the bias indices of adaptation to Chinese numerals are comparable to that in the control condition (adaptation to white noise) at both the single participant level (***Supplementary Fig. 1 Left***, response proportions of adaptation to white noise: 50% for 6, 50% for 8 and 0% for other numbers) and the population level (***Supplementary Fig. 1 Right***, bias indices of adaptation to white noise, *mean* = 0.22 ± 0.38 [95% *CI*]).

These results suggest that semantic processing mechanisms do not take a major role in the perceptual bistability of the partially occluded digital numeral.

## Discussion

In this study, we demonstrate that partial occlusion of digital numerals can induce bistability in their semantic interpretation. Adaptation to visual stimuli that share shape similarities with occluded digital numerals reduces perceptual ambiguities and leads to unique semantic recognition. Our results indicate that the bistable perception of the occluded digital numerals relies neither on early-stage visual processing nor on semantic recognition mechanisms (***Fig. 4***). Rather, it largely depends on the mid-level visual processing stages, which encode complex shapes and symbolic number forms (Conrad et al 2023, Shum et al 2013, Yeo et al 2020, Yeo et al 2017) in a manner invariant to stimulus positions (Ito et al 1995, Logothetis & Sheinberg 1996, Vogels & Orban 1996).

**Figure 4.**
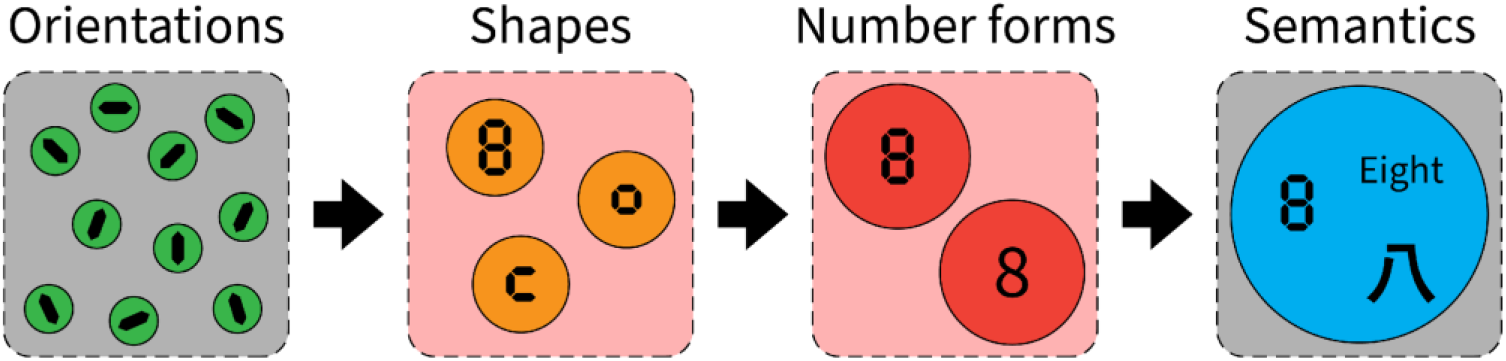
Illustration of the hierarchical visual information processing that endows us with capabilities ranging from the perception of simple orientations to the sensation of complex shapes and number forms, and finally to the semantic understanding of symbols, words, and characters. Our results from visual adaptation implicate the function of middle stages (marked by red squares) in the perceptual bistability of occluded digital numbers.

### Occlusion generates perceptual bistability

In a natural environment, humans often encounter situations where objects of interest are partially covered (***Fig. 1A right***). Nevertheless, recognition of those partially occluded objects takes little effort for the visual system despite the incomplete information. This phenomenon of visual completion has been intensively studied in the fields of psychophysics (Nakayama et al 1989, Shimojo & Nakayama 1990, Tse 1999), physiology (Albright & Stoner 2002), and neuroimaging (Ban et al 2013, Thielen et al 2019). Usually, partial occlusion results in perceptual filling-in of the visual information made unavailable by the occluder and the literature has mainly focused on studying the visual completion of occluded objects (Sekuler & Palmer 1992). However, not all cases of occlusion induce visual filling-in; in some cases, the covered area may be interpreted as containing no relevant information.

Our partially occluded digital numerals can be considered one of such cases since both interpretations are equally probable and the digital numerals are meaningful whether or not the occluded part is filled in. For example, in our experiment, the partially occluded numerals (***Fig. 2A Right***) may be interpreted as either number 6 or 8, depending on whether the area behind the white disk is filled in with a vertical stick. One possible reason for this specific phenomenon is that numerals, such as 6 and 8, are well-learned symbolic characters, and robust visual shape templates exist inside human brains. When occurrences of two numerals (e.g. 6 and 8) are equally probable, the perceptual system must resolve competition between these two perceptual interpretations as occluded digital numerals are processed based on perceptual templates, and such competition causes bistable perception.

We note that there was a considerably large individual difference in the control condition (adaptation to white noise; see ***Supplementary Fig. 1 Right***) . Specifically, some participants consistently reported that they perceived the occluded digital numeral as 6, or consistently as 8, while all participants showed the same trend of perceptual bias after adaptation to digital numerals (***Fig. 3***). The reason why this large individual difference was observed in the control condition is an open question.

### Visual adaptation breaks down the bistable state of partially occluded numerals

In our experiment, the adaptation to numerals and shapes of similar appearances broke down the bistable state and led to a unique perceptual interpretation of the partially occluded digital numerical character. One possible explanation is that adaptation suppresses the sensitivity of the neural mechanism that encodes a certain numerical form. To be specific, the response state of mechanisms encoding numerals 6 and 8 are normally balanced with each other, contributing similarly to the interpretation of a partially occluded numeral and therefore producing perceptual bistability. However, adaptation to the digital numeral 6 led to decreased sensitivity of the corresponding encoding mechanism, resulting in a higher response state of the neural mechanism that encodes digital numeral 8. This process produces a perceptual bias of the partially occluded digital numeral toward the number 8, and vice versa.

## Conclusions

Our findings lead us to conclude that the perceptual bistability of partially occluded digital numerals mainly originates from competitions in visual processing stages involved with processing global shapes and number forms. Neither early visual or semantic processing is likely to be involved in the interpretation of bistable digital numerals.

## Acknowledgments

We thank Kumiko Kobayashi in support of participant recruitment and data curation. We also thank Lothar Spillman and Maiko Uesaki for their suggestions for an earlier version of the manuscript.

## Supplementary materials

**Supplementary Video 1**. The traffic countdown timer is partially covered by the protective roof. Although the digital numerals (from 9 to 5) are temporally aligned in a descending order, they can also be perceptually interpreted as aligned in an ascending order from 5 to 9.

**Supplementary Video 2**. A demo video showing the effect of visual adaptation. Digital numerals of 6 and 8 are presented on the left and right visual fields, respectively. Two occluded digital numerical characters of the same shape are shown on both visual fields after adaptation. Although the test stimuli don’t have any difference, the occluded character on the left visual field looks more like 8, and the other one on the right side looks more like 6.

**Supplementary Video 3**. A demo video showing the stimuli presentation. The adaptation stimulus of the digital numeral 6 on the right visual field is presented for 30 seconds, after that the occluded test stimulus is shown for 0.3 seconds.

**Supplementary Video 4**. A demo video showing the stimuli presentation. The adaptation stimulus of the digital numeral 8 on the right visual field is presented for 30 seconds, after that the occluded test stimulus is shown for 0.3 seconds.

**Supplementary Figure 1.**
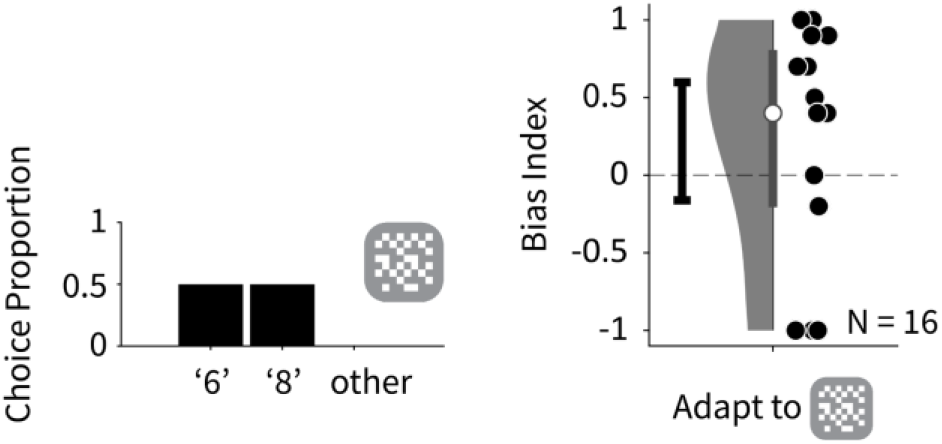
The single participant and populational results of adaptation to white noise control stimulus. Comparing with adapting to character 六 in **Fig. 3D right**, *d* = 0.10, *p* = 1.00, *BF*_*10*_ = 0.29; comparing with adapting to character 八 in **Fig. 3D right**, *d* = 0.04, *p* = 1.00, *BF*_*10*_ = 0.30. Please note that some data points in the right panel are distributed near the upper and lower limits of the figure, indicating large individual differences in bistable perception of the occluded digital numeral in this control condition.

## Notes

### Competing Interest Statement

The authors have declared no competing interest.

